# Anti-inflammatory and pro-proliferative effects of fasudil in human trisomy 21 neural progenitor cells

**DOI:** 10.64898/2026.03.19.712922

**Authors:** Laura L. Baxter, Sarah E. Lee, Kevin A. Fuentes, Iman A. Mosley, Jonathan D. Raymond, Faycal Guedj, Di Zhou, Donna K. Slonim, Elliot J. Glotfelty, David Tweedie, Nigel H. Grieg, Diana W. Bianchi

## Abstract

Down syndrome (DS) results from trisomy for human chromosome 21 and is the most frequent genetic cause of intellectual disability. No effective treatments currently exist that improve neurodevelopment and cognition. Atypical brain development in individuals with DS is apparent before birth, which suggests that the optimal time to begin administration of therapies is prenatally. Human neural progenitor cell (NPC) cultures provide a tractable *in vitro* model system to examine the effects of trisomy 21 (T21) on neurodevelopment and to measure the effects of pharmacological interventions. Here we report the results of preclinical studies evaluating 24 candidate therapies. RNA-Seq analyses found that euploid and T21 NPCs showed different transcriptomic responses to five candidate pharmacotherapies. The Rho-associated coiled-coil kinase (ROCK) inhibitor fasudil increased proliferation of T21 NPCs, reduced expression of inflammatory pathway genes in T21 NPCs, and reduced markers of inflammation in LPS-stimulated microglia model systems. These results demonstrate that fasudil can alter multiple T21-associated abnormalities in a beneficial manner, suggesting that fasudil warrants further study as a candidate prenatal pharmacotherapy for DS.

## Introduction

Down syndrome (DS) is caused by trisomy for human chromosome 21 (trisomy 21; T21) and is the most common genetic cause of intellectual disability. T21 is associated with atypical brain development that is apparent before birth. Magnetic resonance imaging (MRI) and transvaginal neurosonography in living fetuses with T21 have shown that structural alterations and regional brain hypoplasia are apparent in the developing fetal brain (Kitano et al., 2023; Pooh et al., 2025; Tarui et al., 2020; Yun et al., 2021). The early appearance of these neurodevelopmental abnormalities suggests that prenatal treatment may be required to counteract these effects. We and others hypothesize that safe and efficacious prenatal treatment will result in more typical fetal brain growth and development that could lead to improved learning, memory, and independent life skills for individuals with T21 (Guedj et al., 2014; Stagni & Bartesaghi, 2022; Stagni, Giacomini, Guidi, Ciani, & Bartesaghi, 2015)

Prenatal screening for DS provides a unique opportunity for fetal treatment for several reasons. Biochemical, cell-free DNA, and sonographic screening for T21 are part of routine prenatal care in most developed countries, allowing identification of high-risk fetuses during early, critical stages of brain development. Subsequent diagnostic testing via amniocentesis or chorionic villus sampling, followed by karyotype or chromosome microarray, would limit treatment to only those fetuses with confirmed T21. Further, imaging measurements of T21-associated fetal brain changes have already provided baseline phenotypic data that could be used to measure the impact of antenatal therapy on brain growth and development (Kitano et al., 2023; Pooh et al., 2025; Tarui et al., 2020; Yun et al., 2021).

Given the limited availability of fetal brain tissue from individuals with T21, mouse models have previously been used in multiple studies to examine the effects of candidate therapeutics on neurodevelopment during embryonic and neonatal stages that correspond to the human fetal time period. Collectively, these prior efforts examined 15 prenatally- and 12 neonatally-administered molecules (21 molecules total) in the Ts65Dn, Dp(16)1Yey, and Ts1Cje mouse models of DS (Lopez-Hidalgo et al., 2024; Stagni C Bartesaghi, 2022). These studies demonstrated that central nervous system (CNS) development and subsequent cognitive abilities can be improved by pharmacological intervention early in development, with the most significant results arising from prenatal administration (Stagni & Bartesaghi, 2022).

The candidate therapeutics analyzed in prior mouse studies were chosen using a variety of criteria. These included targeting CNS pathways known to be altered in humans with T21 or mouse models of DS, such as the serotonin re-uptake inhibitor fluoxetine (Guidi et al., 2014) or the tropomyosin-related kinase receptor B agonist 7,8-dihydroxyflavone (Giacomini et al., 2019; Stagni et al., 2017; Stagni et al., 2021; Valenti et al., 2021), or inhibiting proteins encoded by trisomic genes such as the antioxidant (-)- epigallocatechin-3-gallate (EGCG), which inhibits DYRK1A (McElyea et al., 2016; Souchet et al., 2019; Stagni et al., 2016; Tielemans et al., 2025). In addition, our laboratory used a transcriptomics approach to identify novel therapeutic candidate molecules (Guedj et al., 2016). Dysregulated gene sets from nine cell and tissue types from human T21 sources and mouse models of DS were used to query the Connectivity Map database (Lamb et al., 2006) to discover molecules predicted to improve dysregulated gene expression. This identified the naturally occurring flavone apigenin (4’,5,7-trihydroxyflavone) as a candidate (Guedj et al., 2016). Subsequent studies (Guedj et al., 2020) showed apigenin reduced oxidative stress and enhanced antioxidant defense responses in T21 amniocytes. In addition, administration of apigenin to pregnant Ts1Cje dams followed by administration to offspring over their lifetimes reduced inflammatory markers, improved neonatal developmental milestones, and improved exploratory behavior and hippocampal long-term memory in adult male mice (Guedj et al., 2020). These results showed that a transcriptomic approach could successfully identify new therapeutic molecules that improve trisomy-associated neurodevelopmental alterations.

Although many candidate molecules showed promise in mouse model studies, similarly beneficial effects were not seen in clinical trials of individuals with DS (Rueda, Florez, et al., 2020). This lack of translation may be due to differences in murine and human neurodevelopment, resulting in different phenotypic effects of trisomy. It may also be attributable to incomplete modeling of trisomy: the Ts65Dn, Dp(16)1Yey, and Ts1Cje mouse models used for these studies are only trisomic for a subset of genes that are orthologous to human chromosome 21 (*Hsa*21). Furthermore, Ts65Dn and Ts1Cje contain aneuploid regions that are not syntenic to *Hsa*21 that could influence therapeutic responses (Davisson et al., 1990; Duchon et al., 2011; Guedj et al., 2023; Reeves et al., 1995; Sago et al., 1998). Overall, these inconsistent results suggest that preclinical models need to be improved to better predict the effects of candidate therapeutics.

The recent creation of human induced pluripotent stem cells (iPSCs) from T21 somatic cells has provided a valuable alternative to mouse models for studying neurodevelopment in DS. These cells are fully trisomic for *Hsa*21 and can be differentiated into a variety of neuronal cell types (Klein & Haydar, 2022; Russo et al., 2024; Watson & Meharena, 2023). We previously created a panel of age- and sex-matched euploid (Eup) and T21 iPSCs and differentiated these iPSCs into neural progenitor cells (NPCs) (Lee et al., 2025). Characterization of these NPCs showed that T21 is associated with genome-wide disruption of the transcriptome, reduced growth, increased oxidative stress, and inter-individual variability in gene and protein expression. NPCs are relevant to T21 neurodevelopmental phenotypes because prenatal brain hypotrophy is associated with fewer proliferating NPCs and reduced neurogenesis (Contestabile et al., 2007; Guidi et al., 2008; Guidi et al., 2018; Lu et al., 2012; Stagni et al., 2018; Stagni et al., 2019, 2020). A previous study showed that the senolytics dasatinib and quercetin together reduced T21-associated cellular phenotypes in NPCs (Meharena et al., 2022), highlighting the utility of NPCs as an *in vitro* model system to test pharmacotherapies.

Here we describe preclinical studies using our panel of T21 NPCs to evaluate candidate therapeutic molecules. Molecules were chosen for analysis based upon previously published literature that suggested their therapeutic potential, or via a transcriptomic approach in which the Library of Integrated Network-Based Cellular Signatures (LINCS) was queried for molecules predicted to reverse T21 NPC gene expression signatures and thus partially restore gene expression to Eup levels (Keenan et al., 2018; Sapashnik et al., 2023). Therapeutic endpoints included proliferation of NPCs, normalization of differentially expressed genes and pathways in T21 NPCs, and reduction of lipopolysaccharide-induced inflammation in mouse macrophages and microglia. These studies found that T21 and Eup cells do not show identical transcriptomic responses to molecules. In addition, these studies demonstrated that the Rho-associated coiled-coil kinase (ROCK) inhibitor fasudil increased proliferation in T21 NPCs, reduced expression of inflammatory pathways in T21 NPCs, and showed anti-inflammatory properties in Eup microglia, suggesting fasudil shows potential for prenatal pharmacotherapy for DS.

## Results

### Selection of candidate therapeutic molecules

Extending upon our previous transcriptomics approach to identify novel therapeutic candidate molecules (Guedj et al., 2016), a set of differentially expressed genes (DEGs) in T21 NPCs relative to Eup NPCs (Lee et al., 2025) was used to query LINCS (https://lincsproject.org/). LINCS is an NIH-funded, publicly available data repository of information on the cellular responses of over 70 human cell types to various genetic, disease, or chemical perturbations, including transcriptional responses to over 20,000 bioactive small molecules (Keenan et al., 2018). Submission of gene signatures for the T21 DEGs (DEGs plus expression changes for each gene) to LINCS identified 165 molecules with weighted connectivity scores < -0.2 in NPCs (Subramanian et al., 2017)(Fig. 1A). Negative scores indicate reversal of the gene signature query, therefore these 165 molecules were predicted to correct gene expression in T21 NPCs towards Eup levels.

**Figure 1.**
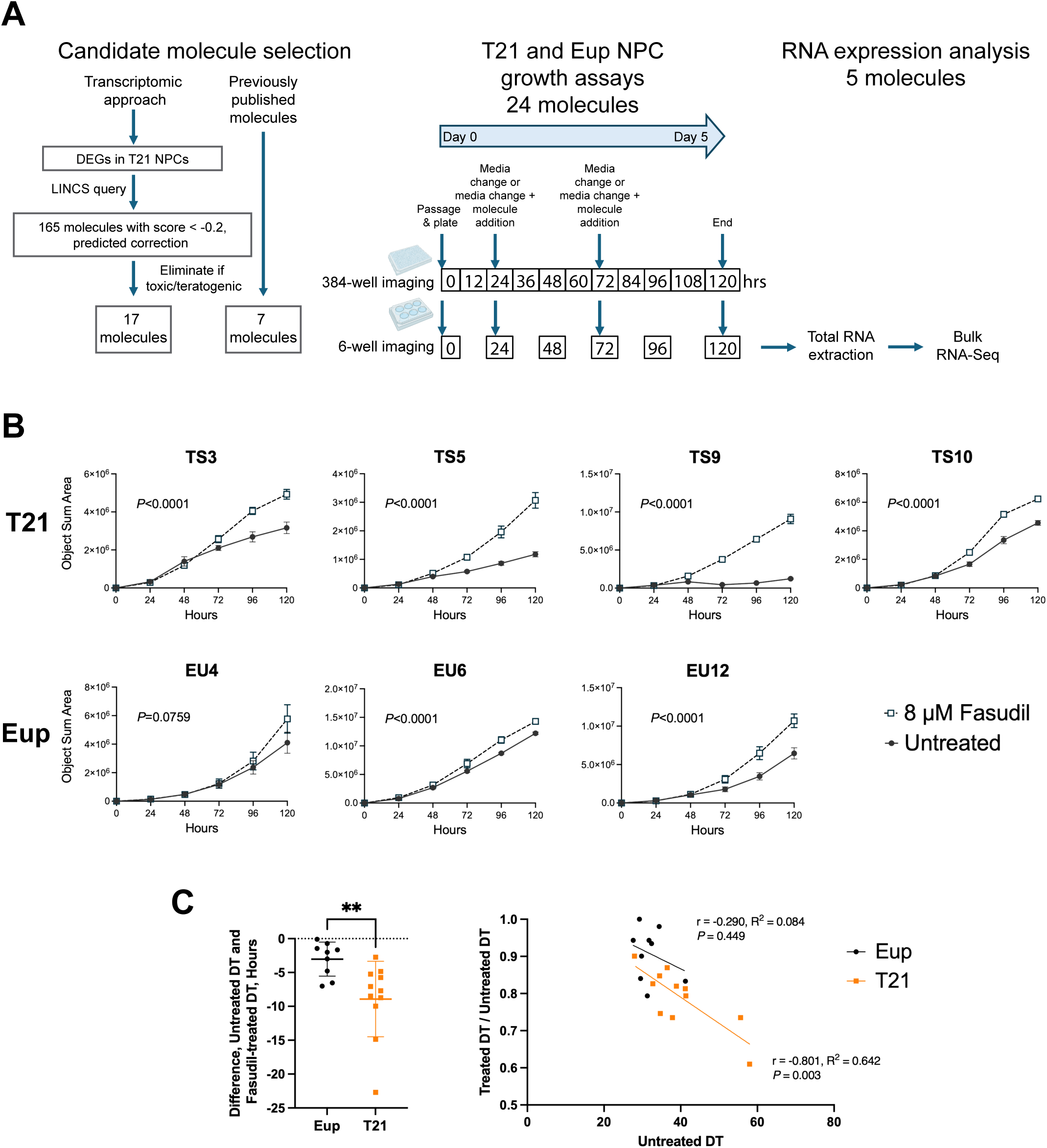
Analyses of a panel of candidate therapeutics showed that fasudil improved T21 NPC proliferation. A) Experimental design. Twenty-four candidate molecules were selected from a transcriptomic approach, using di=erentially expressed genes (DEGs) in T21 NPCs to query LINCS, or from previously published results suggesting therapeutic benefits. Molecules were screened for cell proliferation e=ects in 384-well plate format followed by 6-well plate format by quantitative live-cell imaging every 12 or 24 hrs, respectively. Bulk RNA-Seq analysis was performed on RNA collected from Eup and T21 NPCs treated with five molecules after 120 hrs. B) Increased growth occurred following 8 µM fasudil treatment in all T21 lines and in a subset of Eup lines. Each graph shows a single experiment for one cell line in the 6-well plate format experiments, and each point is the mean of 6 replicate wells. Open squares = fasudil treated, closed circles = untreated. T21 and Eup lines are named using the prefixes TS and EU, respectively, and these lines are described in Supp. Tab. S2. C) Greater reductions in doubling time (DT) occurred in T21 NPCs (graph, left). Mean DT changes after fasudil treatment = -8.92 hrs (SD=5.6) for T21 and -3.02 hrs (SD=2.5) for Eup (*P* = 0.0016, Mann-Whitney two-tailed test). Fasudil-treated doubling time normalized to untreated doubling time showed greater e=ects in T21 NPCs (graph, right). The ratio of treated/untreated DT was significantly correlated with untreated DT in T21 lines, consistent with fasudil showing greater e=ects in slower growing lines, while Eup NPCs showed no correlation. N = 11 for T21, 9 for Eup; each point shows DT for one experiment for a single NPC line (data in Supp. Tab. S1c). Black = Eup lines, orange = T21 lines.

Seventeen of these molecules with no reported teratogenicity or contraindications in pregnancy were chosen for further examination using NPC cultures (Fig. 1A, Supp. Tab. S1a). Of note, previous studies on seven of these transcriptomically-selected molecules showed beneficial effects in preclinical models of T21, neurodevelopment, neurodegeneration, or oxidative stress. These included: the fatty acids alpha linolenic acid (ALA) and eicosapentaenoic acid (EPA) (Garcia-Cerro et al., 2020; Martinez-Cue & Bartesaghi, 2022; Zmijewski et al., 2015), the PGC-1α activator metformin (Izzo et al., 2017), the flavonoids naringenin and nobiletin (Ghasemi-Tarie et al., 2022; Jahan et al., 2024; Nakajima & Ohizumi, 2019; Nouri et al., 2019; Xiong et al., 2023), the PPAR-γ agonist pioglitazone (Mollo et al., 2019), and the lipid sphingosine (Hwang et al., 2019; Tang et al., 2017). No information on therapeutic effects in preclinical models of T21 was found in PubMed for the remaining 10 molecules identified by the LINCS query (chloroquine, cilnidipine, geranylgeraniol, glycodeoxycholic acid, imperatorin, isoxsuprine, pinocembrin, tangeritin, trapidil, and vitexin).

Alongside the candidate therapeutic molecules identified by transcriptomic analysis, seven molecules with previously published data suggesting potential therapeutic benefits in T21 model systems or individuals with DS were also chosen for analysis in NPCs (Fig. 1A, Supp. Tab. S1a). These molecules were apigenin, which we previously identified using a transcriptional approach (Guedj et al., 2020), the flavonoid 7,8 dihydroxyflavone (Giacomini et al., 2019; Stagni et al., 2017; Stagni et al., 2021; Valenti et al., 2021), choline chloride (Alldred et al., 2023; Alldred et al., 2025; Gautier et al., 2023; Gautier et al., 2024; Moon et al., 2010; Powers et al., 2017; Powers et al., 2021; Velazquez et al., 2013), the antioxidants curcumin (Rueda, Vidal, et al., 2020) and EGCG (Blazek et al., 2015; Catuara-Solarz et al., 2015; de la Torre et al., 2016; De la Torre et al., 2014; McElyea et al., 2016; Souchet et al., 2019; Stagni et al., 2016; Starbuck et al., 2021; Tielemans et al., 2025), the ROCK inhibitor fasudil (LeBlanc-Straceski et al., 2022; Lopez-Hidalgo et al., 2024), and the selective serotonin-reuptake inhibitor fluoxetine (Bianchi et al., 2010; Guidi et al., 2014; Guidi et al., 2013; Heinen et al., 2012; Stagni, Giacomini, Guidi, Ciani, Ragazzi, et al., 2015; Stagni et al., 2013).

### Analysis of cell proliferation in response to candidate therapeutic treatment

T21 NPCs show reduced proliferation (Contestabile et al., 2007; Guidi et al., 2008; Guidi et al., 2018; Lee et al., 2025; Lu et al., 2012; Stagni et al., 2018; Stagni et al., 2019, 2020), thus we examined the effects of candidate pharmacotherapies on NPC growth. These experiments used a panel of NPCs derived from iPSC lines that were previously generated from fibroblasts of unrelated Eup individuals and individuals with T21 (Lee et al., 2025). First, 17 iPSC lines (nine Eup and eight T21, Supp. Tab. S2) were stably transduced with Nuclight Red, a nuclear-restricted, non-integrating, lentivirus-derived fluorescent vector. Transduced iPSCs were then differentiated into NPCs that retained Nuclight Red expression, which permitted automated counting of live NPCs for measurement of growth.

T21 NPC lines exhibited variable recovery from passaging when supplemented with the commonly used ROCK inhibitor Y-27632. To address this issue, we tested a protocol during neural induction from iPSCs that reportedly improved survival during passaging by supplementation of media with the small molecule “CEPT” cocktail of <u>c</u>hroman 1, <u>e</u>mricasan, <u>p</u>olyamines, and trans-ISRIB (Chen et al., 2021). Administration of CEPT during passaging of three Eup and three T21 lines led to greater cell survival following passaging in both genotypes (Supp. Fig. S1). Therefore, CEPT supplementation was used for passaging throughout all NPC analyses.

A 384-well plate format was used to examine the effects of candidate pharmacotherapies on NPC growth (Fig. 1A). A panel of Eup and T21 Nuclight Red-transduced NPC lines (4-5 Eup and 5-6 T21 lines per molecule, Supp. Tab. S2) was tested for each molecule. Eup and T21 NPCs were cultured for 120 hrs in the presence or absence of candidate molecules at two or three concentrations (based on previous *in vitro* dose range-finding applications), and quantitative live-cell imaging was performed every 12 hrs to generate growth curves. Each NPC line was evaluated in quadruplicate wells for each molecule, and growth assays were repeated for every molecule. Eight molecules caused decreased growth and increased doubling time (DT) in Eup and T21 NPCs relative to untreated NPCs, indicating potential toxic effects (Supp. Tab. S1b). The remaining molecules showed no change in DT or variable, inconsistent effects on DT. Of note, 8 µM fasudil showed improved growth and reduced DT in 18% of the experimental replicates across all T21 cell lines, the most of any molecules tested in the 384-well plate format (Supp. Tab. S1b).

### Fasudil consistently increases cell growth in T21 NPCs

Additional analyses of the effects of candidate therapeutics on cell proliferation were performed on NPCs in 6-well plates (Fig. 1A), to determine if similar growth effects were seen in large volume culture conditions and to generate samples for RNA collection. Five molecules were used: two previously identified candidate therapeutics, fasudil (8 µM) and EGCG (8 µM), and three transcriptionally identified molecules, apigenin (4 µM), isoxsuprine (1 µM), and sphingosine (2 µM). EGCG, apigenin, and isoxsuprine did not consistently change growth (Supp. Tab. S1c). Sphingosine showed no consistent effects on growth in T21 NPCs but consistently increased DT (thus slowing growth) in two Eup lines (Supp. Tab. S1c). Interestingly, fasudil reduced DT (thus improving growth) in four T21 NPC lines, and this effect was reproducible across all replicate experiments (Supp. Tab. S1c).

Two-way ANOVA analyses of growth curves confirmed significantly increased growth in all fasudil-treated T21 NPCs across all replicate experiments (Fig. 1B, Supp. Tab S1d). In Eup lines, fasudil showed less consistent effects on growth in comparison to T21 lines. Fasudil consistently reduced DT in Eup line EU12, but Eup lines EU4 and EU6 showed reduced DT in only one or two experiments (Supp. Tab. S1c). In two-way ANOVA analyses of Eup growth curves, EU4 showed no change, EU6 showed consistently increased growth across all three experiments, and EU12 showed increased growth in two out of three experiments (Fig. 1B, Supp. Tab. S1d).

Several T21 lines showed slower growing times / longer DTs than Eup lines in untreated conditions (Supp. Tab. S1c). Therefore, we wanted to determine if fasudil caused stronger growth effects in T21 NPCs. Indeed, greater reductions in DT were seen in T21 NPCs in comparison to Eup NPCs in the 6-well format cultures, with mean DT changes after fasudil treatment of -8.92 hrs (SD=5.6) for T21 and -3.02 hrs (SD=2.5) for Eup (*P* = 0.0016, Mann-Whitney two-tailed test, Fig. 1C). T21 lines showed greater variability in DTs, and the largest change in DT was seen in the slowest growing T21 line (line TS5, mean DT change = -14.27 hrs (SD=8.7)). The ratio of treated/untreated DT was correlated with untreated DT in T21 lines (r = -0.801, R^2^ = 0.642, *P* = 0.003, Fig. 1C), consistent with fasudil showing greater effects in slower growing lines. In contrast, no correlation was present between the ratio of treated/untreated DT and untreated DT in Eup NPCs (Fig. 1C).

### Anti-inflammatory properties of fasudil, apigenin, and isoxsuprine

Candidate pharmacotherapies with anti-inflammatory properties are of interest because upregulated interferon and inflammatory pathways contribute to multiple phenotypes in mouse models of DS and individuals with T21 (Chi et al., 2023; Galbraith et al., 2023; Rachubinski et al., 2024; Sullivan et al., 2017; Sullivan et al., 2016; Tuttle et al., 2020; Waugh et al., 2023). Microglia are resident macrophages in the brain that function as the brain’s innate immune system. As a source of inflammatory signaling, microglia play integral roles in shaping neural circuits and maintaining brain homeostasis, which is crucial during development of the fetal brain (Anderson et al., 2025; Lawrence et al., 2024). *In vitro* microglia/macrophage cultures have previously been used to recapitulate *in vivo* and primary microglial inflammatory phenotypes across many drug classes (Glotfelty et al., 2023; Hsueh et al., 2025; Kopp et al., 2024; Lecca et al., 2022; Li et al., 2021). These culture systems were applied to the candidate molecules fasudil, EGCG, apigenin, isoxsuprine, and sphingosine using the classical lipopolysaccharide (LPS) challenge model, in which LPS induces an inflammatory response that is mediated by Toll-like receptor 4.

LPS challenge was first applied to the RAW 264.7 cell line, a mouse macrophage cell line reported to have parallel features to blood-induced microglia (Cheon et al., 2017; Mendiola et al., 2023). Cellular viability evaluation at 1, 10, and 100 µM concentrations showed that EGCG, fasudil, apigenin, and sphingosine were toxic to RAW 264.7 cells at 100 µM concentrations, reducing the levels of cell viability to 80±2%, 33±1%, 55±1%, and < 1% compared to the control group (vehicle), respectively. Consequently, the highest compound concentration assessed in subsequent studies was 30 µM.

In LPS-stimulated RAW 264.7 cells, fasudil, apigenin, and isoxsuprine all showed strong anti-inflammatory effects, evidenced by significant reductions in media levels of nitrite, TNF-α and IL-6 (Fig. 2, Supp. Tab. S3a). Fasudil at 30 µM reduced the levels of nitrite, IL-6 and TNF-α with no cell toxicity, and fasudil at 10 µM also showed reduced IL-6 levels. Apigenin potently reduced nitrite, IL-6 and TNF-α at 30 µM; however, at this concentration there was evidence of a mild but statistically significant level of cell toxicity.

**Figure 2.**
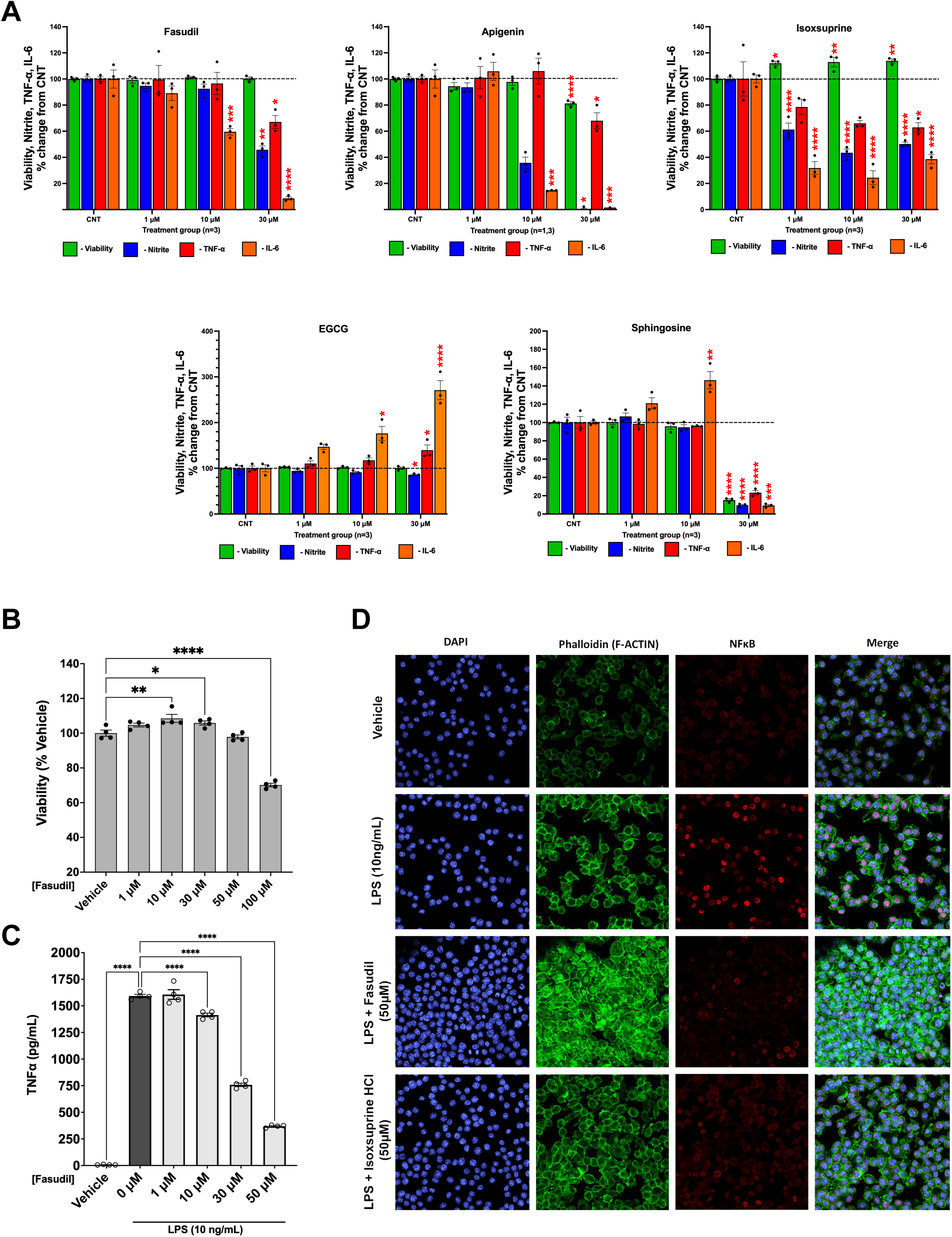
Effects of fasudil, apigenin, isoxsuprine, EGCG, and sphingosine in LPS- challenged RAW 264.7 cells and IMG cells. A) Viability effects and levels of nitrite, TNF-α, and IL-6 in RAW 264.7 cells. Fasudil, apigenin, and isoxsuprine showed strong anti-inflammatory effects, evidenced by reduced media levels of nitrite, TNF-α and IL-6 relative to untreated control (CNT) samples. In contrast, EGCG and sphingosine showed moderate pro-inflammatory effects, evidenced by increased media levels of IL-6 and TNF-α for EGCG. EGCG also showed modest anti-inflammatory activity 30 µM, evidenced by reduced nitrite levels. Toxicity was observed for sphingosine at 30 µM. B) Treatment of IMG cells with fasudil (no LPS) showed that fasudil increased cell viability at 10 µM and 30 µM but was toxic at 100 µM. C) Fasudil induced dose-dependent reductions in secreted TNF-α in LPS-challenged IMG cells, indicating significant anti-inflammatory effects. D) LPS-challenged IMG cells showed increased nuclear expression of NFkB, and fasudil and isoxsuprine at 50 µM both reduced nuclear localization of NFkB (in red). All data shown as mean ±SEM. Statistical significance was determined by ordinary one-way ANOVA with Dunnett’s multiple comparisons test or Kruskal-Wallis test with Dunn’s multiple comparisons test, with significance indicated as follows: * p<0.05, ** p<0.01, *** p<0.001, **** p <0.0001.

Apigenin at 10 µM reduced levels of nitrite and IL-6 with no cell toxicity. Isoxsuprine strongly reduced all markers of inflammation at all 3 concentrations with no cellular toxicity (high variability limited the significance of TNF-α reduction at 1 µM and 30 µM) and also increased viability.

In contrast, EGCG showed both pro- and anti-inflammatory effects and sphingosine showed modest pro-inflammatory effects in LPS-stimulated RAW 264.7 cells (Fig. 2, Supp. Tab. S3). Both compounds generated elevations in IL-6 in a concentration-dependent manner. These were statistically significant for EGCG at 10 and 30 µM and for sphingosine at 10 µM. Sphingosine at 30 µM proved to be toxic to RAW 264.7 cells (15±1% viability compared to control). Only EGCG at 30 µM was able to reduce nitrite levels with no cellular toxicity.

The anti-inflammatory effects of fasudil and isoxsuprine were subsequently validated in LPS-challenged mouse microglial IMG cells. This immortalized microglial line closely models brain function, is widely used in studies of brain disease (Banerjee et al., 2021), and exhibits potent anti-inflammatory effects from ROCK and pan-kinase inhibitors (Glotfelty et al., 2023). Viability testing of IMG cells treated with fasudil from 1 to 100 µM demonstrated moderate but significant elevations in cell viability up to a concentration of 30 µM (Fig. 2B, Supp. Tab. S3b), which paralleled the increased growth seen following fasudil treatment in NPCs (Fig. 1B). Fasudil at 100 µM demonstrated cellular toxicity, thus a maximal concentration of 50 µM was used in subsequent LPS studies. Viability testing in LPS-challenged IMG cells confirmed that fasudil remained non-toxic from 1 to 50 µM (data not shown).

Fasudil reduced the levels of secreted TNF-α in LPS-challenged IMG cells in a concentration-dependent manner (Fig. 2C, Supp. Tab. S3b). Furthermore, immunofluorescence showed that while LPS stimulation of IMG cells caused elevated nuclear expression of NFkB, indicating that NFkB has been released from cytoplasmic sequestration and has entered the nucleus to activate target genes, treatment of IMG cells with fasudil or isoxsuprine prior to LPS stimulation reduced NFkB nuclear staining (Fig. 2D). These results show that both fasudil and isoxsuprine dampen the NFkB pathway, a novel finding related to isoxsuprine and consistent with previous findings related to fasudil in other disease models and cell types (Guo et al., 2020; Hou et al., 2012; Liu et al., 2022; Okamoto et al., 2010; Song et al., 2013).

### Fasudil partially corrects dysregulated gene expression and pathways in T21 NPCs

To examine the effects of fasudil treatment on Eup and T21 NPC RNA expression, bulk RNA-Seq analysis was performed. Total RNA for RNA-Seq was collected at 120 hrs from Eup and T21 NPC lines (N = 3 and 4, respectively) that were cultured in the 6-well plate format in the presence or absence of fasudil (Fig. 1A). Multidimensional scaling (MDS) of all RNA-Seq data showed separation of Eup and T21 NPC lines (Fig. 3A). T21 NPC lines showed more variability than Eup lines by MDS, corresponding with the variability previously observed for these lines by microarray and proteomics analyses (Lee et al., 2025). Genotype comparison of untreated T21 (T21Unt) and untreated Eup (EupUnt) samples identified 8466 DEGs (T21 DEGs, *P*-Adj ≤ 0.05, FC ≥|1.5|), reflecting the widespread gene expression changes that occur from T21 (Supp. Tab. S4a). Ingenuity pathway analysis (IPA) showed that enriched pathways associated with T21 DEGs included upregulation of cellular stress and injury, cellular immune response, and immune system pathways along with downregulation of cell cycle pathways (Fig. 3B, Supp. Tab. S4b), matching previous analyses of microarray data from these NPC lines (Lee et al., 2025).

**Figure 3.**
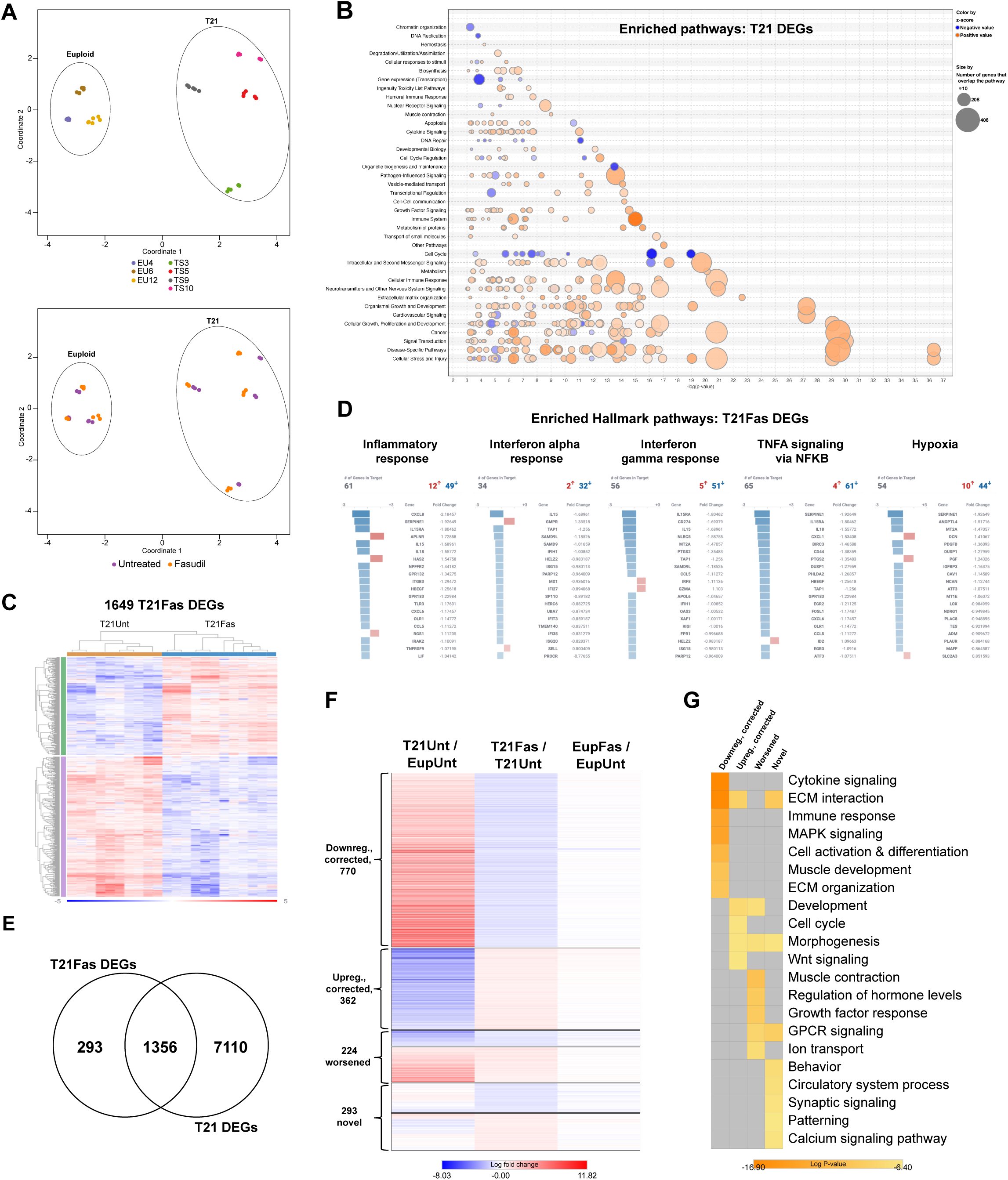
RNA-Seq analysis of fasudil-treated NPCs showed that fasudil partially corrected dysregulated gene expression and immune system pathways in T21 NPCs. A) MDS plot of all RNA-Seq data for T21 and Eup NPCs treated with fasudil shows strong separation of NPC cell lines by genotype and moderate separation of fasudil-treated samples from untreated samples, with greater separation seen in T21. Top plot is colored by individual NPC lines to show genotype separation, and bottom plot is colored by fasudil-treated samples (orange) and untreated samples (purple). T21 and Eup lines are named with the prefixes TS and EU, respectively. B) Ingenuity Pathway Analysis (IPA) showed top pathway categories of significantly enriched pathways for T21Unt vs EupUnt NPCs, termed T21 DEGs. Blue = negative Z score / pathway downregulation, and orange = positive Z score / pathway upregulation. Circle size represents the number of DEGs in the pathway. Upregulation of immune system pathways and downregulation of cell cycle pathways are present. C) Hierarchical clustering of 1649 DEGs identified between T21 fasudil-treated NPCs (T21Fas) and T21 untreated NPCs (T21Unt), termed T21Fas DEGs. D) Heat maps of the top 20 genes (by |FC|) in significantly enriched MSigDB Hallmark pathways, illustrating downregulation of inflammatory response, interferon alpha and gamma responses, TNF-α signaling by NFkB, and hypoxia pathways in T21Fas DEGs. E) Venn diagram showing 1356 DEG overlap between T21Fas DEGs and T21 DEGs. F) Heatmap of log fold-change values for the 1649 T21Fas DEGs in T21Unt vs. EupUnt (left), T21Fas vs. T21Unt (center), and EupFas vs. EupUnt (right). Heatmap rows are clustered based upon correction of T21 DEGs by T21Fas DEGs (770 downregulated, corrected, and 362 upregulated, corrected), worsening of T21 DEGs by T21Fas DEGs (224 worsened, middle), and T21Fas DEGs that did not overlap with the T21 DEG set (293 novel, bottom). G) Pathway analyses of the four sub-clusters of T21Fas DEGs showed that the most significantly enriched pathways are in the downregulated, corrected DEGs and affect immune system functions.

Examination of MDS clusters for individual T21 NPC lines showed separation of fasudil-treated and untreated samples (Fig. 3A, bottom). This separation was also apparent in two Eup lines, but to a lesser degree, indicating extensive gene expression changes from fasudil treatment in T21 NPCs. Concordant with this result, differential gene expression analysis between fasudil-treated T21 (T21Fas) and T21Unt samples identified 1649 DEGs (*P*-Adj ≤0.05, FC ≥|1.5|, Fig. 3C, Supp. Tab. S5a). Twenty-one of these genes were encoded on *Hsa*21 (Supp. Tab. S5b).

The most significantly enriched MSigDB Hallmark pathways in the 1649 DEGs showed downregulation of Interferon alpha, Interferon gamma, Inflammatory response, and TNF-α signaling by NFkB (Fig. 3D, Supp. Tab. S5c), corresponding with the anti-inflammatory properties observed in RAW 264.7 and IMG cells following fasudil treatment (Fig. 2). The Hallmark Hypoxia pathway was also significantly enriched and downregulated (Fig. 3D, Supp. Tab. S5c), which is notable because increased reactive oxygen species have been previously reported in T21 neurons (Briggs et al., 2013; Busciglio & Yankner, 1995). IPA significantly enriched pathways also included numerous cellular immune response and cellular stress and injury pathways, including downregulation of Interferon signaling, neuroinflammation, and CGAS-STING pathways (Supp. Fig. S2, Supp. Tab. S5d).

To determine if fasudil treatment corrected altered gene expression in T21 cells, the 1649 T21Fas DEGs were compared to the 8466 T21 DEGs, and 1356 overlapping DEGs were identified (Fig. 3E, Supp. Tab. S6a). Interestingly, fasudil partially corrected T21 gene dysregulation in most overlapping DEGs, as 83.5% (1132/1356) of the T21Fas DEGs showed expression changes in the opposite direction from those in T21 DEGs (Fig. 3F, Supp. Tab. S6b), i.e., dysregulated genes in the genotype comparison were attenuated in the T21 fasudil treatment comparison. The 1132 corrected DEGs included 13 *Hsa*21-encoded T21 DEGs (Supp. Tab. S6c). Hierarchical clustering of the fold-change values for these subsets of genes (Fig. 3F) illustrated that ∼2/3 of the 1132 corrected genes were downregulated by fasudil (770 genes). Only 224 T21Fas DEGs (16.5%) worsened T21-associated gene expression changes, and the remaining 293 DEGs (17.8%) were novel (Fig. 3F, Supp. Tab. S6a).

Correction by fasudil was also apparent at the pathway level. Comparison of IPA enriched pathway lists identified 64 T21Unt vs EupUnt pathways (with predicted directionality and IPA Z-score >|2|) that were rescued by fasudil in T21 NPCs (Supp. Tab. S6d). The majority were downregulated by fasudil, including many immune system-related pathways such as interferon alpha/beta signaling, CGAS-STING signaling, interferon gamma signaling, RHO GTPase cycle, and senescence. Rescued pathways that were upregulated by fasudil related to cell cycle and chromosome replication, pathways that are characteristically slowed in T21 cells. Upregulation of Sonic Hedgehog (SHH) signaling was also present, which is intriguing because SHH pathways are impaired in mouse models of DS, and restoration of SHH expression or downstream signaling can alleviate abnormal neurological phenotypes (Das et al., 2013; Gao et al., 2021; Roper et al., 2006).

Metascape pathway analysis on the four DEG subsets – downregulated, corrected; upregulated, corrected; worsened; and novel (Fig. 3F) – showed that the most significantly enriched pathways were in the downregulated, corrected DEG set (Fig. 3G). Two of the top downregulated pathway classes were cytokine signaling and immune response, consistent with fasudil correcting the elevated immune system pathways in T21 NPCs (Fig. 3G). The upregulated, corrected DEGs were enriched for development and cell cycle pathways, corresponding to the increased growth seen with fasudil treatment (Fig. 3G). The worsened DEG pathways had less significant average *P-*values than the downregulated, corrected pathways, and included pathways related to muscle contraction, regulation of hormone levels, and growth factor response. Novel DEGs were enriched for a variety of pathways, including GPCR signaling, behavior, and circulatory system processes. Taken together, these transcriptomic analyses indicate that fasudil causes widespread and primarily beneficial changes in T21 gene expression, with the strongest effects seen in reduction of elevated immune system pathways.

The fasudil-induced gene expression changes in Eup NPCs differed from those in T21 NPCs. Fold-change values for fasudil-treated Eup (EupFas) vs EupUnt NPCs using the 1649 T21Fas DEG set indicated no or minimal expression changes in these genes in Eup NPCs (Fig. 3F, right). Differential gene expression analysis between EupFas and EupUnt NPCs identified only 172 DEGs, which was ∼10-fold fewer than those in T21 lines (Supp Tab. S7a). Unlike T21 NPCs, pathway enrichment analyses for Eup NPCs did not identify downregulation of inflammatory pathways from fasudil treatment. Upregulation of *WNT1, WNT10B, WNT2B, WNT3A,* and *WNT4* expression was present in fasudil-treated Eup NPCs, leading to enrichment of multiple WNT signaling pathways regulating growth, proliferation, and development (Supp Tab. S7b).

Comparison of the T21Fas and EupFas DEG gene sets (1649 and 172 DEGs, respectively) identified 63 shared DEGs, 52 of which were changed in the same direction by fasudil (Fig. 4A, Supp. Tab. S7c). This shared list included upregulation of genes encoding the transcription factors LMXA and DMBX1, the enzyme DCT, and the signaling molecules BMP4 and WNT4, all of which regulate NPC proliferation and forebrain development. Most of the DEGs that were changed in the same direction (40/52, 77%) showed greater fold-change values in T21 (Supp Tab. S7c), which may reflect the stronger growth effects observed in T21 (Fig. 1C).

**Figure 4.**
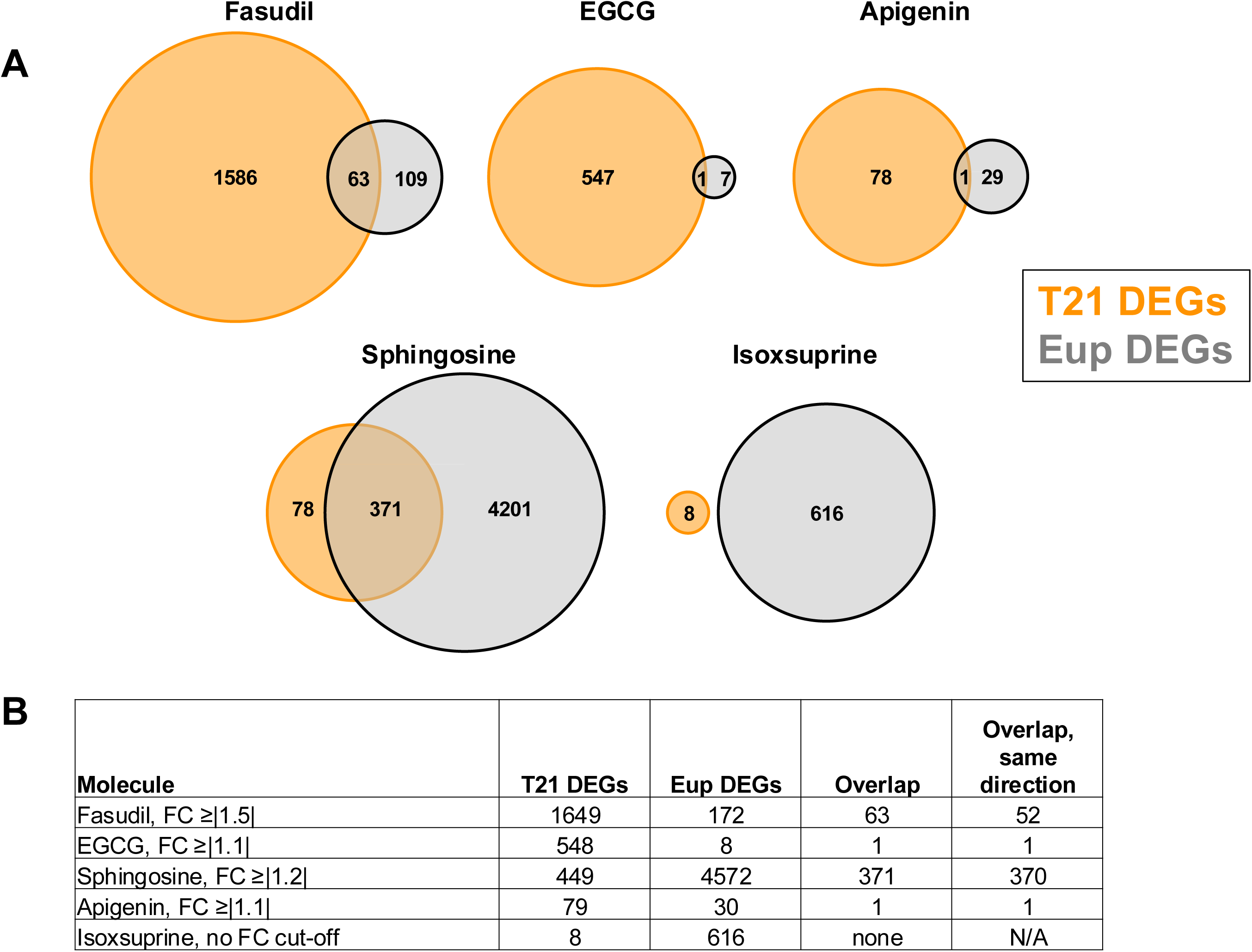
Euploid and T21 NPCs showed different transcriptomic responses to candidate pharmacotherapies. A) Venn diagrams illustrating that T21 NPCs (orange) showed greater numbers of DEGs than Eup NPCs (gray) in response to fasudil, EGCG, and apigenin. In contrast, Eup NPCs showed more DEGs in response to sphingosine and isoxsuprine. B) Table of DEGs for T21, Eup NPCs, DEGs that overlap between T21 and Eup, and overlapping T21 and Eup DEGs that changed in the same direction in response to each candidate molecule. All DEGs used a cut-off of *P*-Adj ≤0.05, and fold-change (FC) cut-off values are shown for each molecule’s DEG set. For isoxsuprine, most FC values were <|1.1|, so no FC cut-off was used.

### Effects of EGCG, apigenin, isoxsuprine and sphingosine on T21 NPCs

Bulk RNA-Seq analyses were also performed on total RNA from Eup and T21 NPCs that were treated with EGCG, apigenin, isoxsuprine, or sphingosine. Differential gene expression analysis was first performed between treated and untreated T21 NPCs for each candidate pharmacotherapy.

In contrast to fasudil’s effects, EGCG showed modest effects on gene expression in T21 NPCs, indicated by only one DEG at p-Adj. ≤0.05, FC ≥|1.5| and 30 DEGs at p-Adj. ≤0.05, FC ≥|1.2|. Therefore, a statistical cut-off of p-Adj. ≤0.05, FC ≥|1.1| was used, giving 548 T21EGCG DEGs (Supp. Tab. S8a). Enriched pathway analysis showed upregulation of multiple pathways related to extracellular matrix organization and cell surface interaction (Supp. Tab. S8b). In addition, most EGCG-induced gene expression changes did not correct T21 dysregulation: 374 T21EGCG DEGs overlapped with the 8175 T21DEGs (untreated cells in the EGCG experiment, Supp. Tab. S8c), and 84.5% of these showed FC values in the same direction (316/374) (Supp. Tab. S8d). Enriched pathway analysis of the overlapping 374 DEG set (Supp. Tab. S8e) showed upregulation of multiple pathways governing cancer/cell growth, extracellular matrix organization, and cellular stress and injury. Several pathways relating to cellular immune response / immune system were also upregulated, which may correspond to the modest pro-inflammatory effects of EGCG that were seen in LPS-stimulated RAW 264.7 cells. Overall, these results indicate EGCG caused modest changes in T21 NPC gene expression, and these changes primarily worsened T21-associated gene dysregulation.

Sphingosine treatment of T21 NPCs also showed more modest changes in gene expression in comparison to fasudil, with fewer DEGs and smaller FCs. In sphingosine-treated T21 NPCs (T21Sph) relative to untreated T21 NPCs (T21Unt), only six DEGs were present using a cut-off of *P*-Adj ≤0.05, FC ≥|1.5|, thus a limit of *P*-Adj ≤0.05, FC ≥|1.2| was applied, which gave 449 DEGs (Supp. Tab. S9a). This gene list included upregulation of multiple genes encoding collagens and matrix metalloproteinases. IPA and Metascape pathway analyses of the 449 DEG set identified upregulation of multiple pathways related to extracellular matrix organization as well as enrichment of pathways regulating synaptic transmission, development, and behavior (Supp. Tab. S9b,c). Approximately half of the T21Sph vs T21Unt DEGs (244/449 DEGs) overlapped with the 5127 T21DEGs (untreated cells in the sphingosine experiment, Supp. Tab. 9d), and 85% of the sphingosine-induced gene expression changes showed FC values in the same direction as the T21DEGs (207/244), thus worsening T21-associated gene expression changes (Supp. Tab. S9e). The shared DEG list included upregulation of *TGFB1*, of note because increased TGFB1 expression occurs in T21 iPSC-derived NPCs, and has been suggested to contribute to the altered balance between neurogenesis and gliogenesis in T21 NPCs (Huang et al., 2025). Enriched pathway analysis of the overlapping 244 DEG set (Supp. Tab. S9f) identified upregulation of pathways related to extracellular matrix organization, pathogen induced cytokine storm signaling, and HIF1 signaling. Overall, these results show no clear improvements in T21 gene dysregulation and indicate sphingosine may exacerbate T21-associated gene dysregulation in NPCs.

Apigenin and isoxsuprine showed minimal effects on gene expression in T21 NPCs. For apigenin, there were no DEGs (P-Adj ≤0.05) with FC values ≥|1.2|, and only 79 DEGs showed FC values ≥|1.1| (Supp. Tab. S10a). Only eight DEGs were present for isoxsuprine, and these DEGs had extremely low FCs (<|1.1|) (Supp. Tab. S10b). The small numbers of genes in the apigenin and isoxsuprine T21 DEG lists prevented enriched pathway analyses.

### Euploid and T21 NPCs show different responses to candidate pharmacotherapies

As mentioned above, T21 NPCs showed more transcriptional changes than Eup NPCs in response to fasudil. Interestingly, differential gene expression analysis between treated and untreated Eup NPCs for each candidate pharmacotherapy (Supp Tab. S11a-d) revealed that the magnitude of transcriptomic changes for T21 and Eup NPCs also differed for the other four candidate therapeutic molecules (Fig. 4). T21 NPCs showed more DEGs than Eup in response to fasudil, EGCG, and apigenin, and Eup NPCs showed more DEGs than T21 in response to sphingosine and isoxsuprine (Fig. 4A, B). Furthermore, there was little to no overlap of T21 and Eup DEGs in EGCG, apigenin, or isoxsuprine, indicating distinct transcriptional effects by genotype. Although there was greater overlap of DEGs in fasudil and sphingosine, and the majority of DEGs changed in the same direction for both T21 and Eup, there were striking differences in the number of DEGs between T21 and Eup for these two molecules, with ∼10-fold more DEGs in T21 NPCs for fasudil and ∼10-fold more DEGs in Eup NPCs for sphingosine. Of note, pathway analyses of the sphingosine Eup DEGs identified significant upregulation of a variety of immune response pathways, including interferon alpha and interferon gamma responses, inflammatory response, and TNF-α signaling by NFkB (Supp Tab. S11e,f), supporting the pro-inflammatory effects of sphingosine in Eup cells seen in RAW 264.7 cells. Taken together, these results suggest that the transcriptomic responses of T21 NPCs to pharmacotherapies may differ from what is expected based upon Eup NPC responses.

## Discussion

This study used *in vitro* cultures of Eup and T21 NPCs to evaluate the ability of candidate pharmacotherapies to alleviate T21-associated cellular phenotypes that may be relevant to atypical neurodevelopment in DS. The ROCK inhibitor fasudil consistently increased proliferation in T21 NPCs, suggesting fasudil could improve the reduced proliferation that is seen in T21 neurodevelopment. Fasudil also demonstrated anti-inflammatory effects, evidenced by reduced RNA expression of inflammatory pathway genes in T21 NPCs and reduced inflammatory markers in LPS-challenged microglia model systems. Previous studies have shown that numerous phenotypes in mouse models of DS and individuals with T21 are associated with upregulated interferon and inflammatory pathways, indicating the relevance of candidate therapeutics with anti-inflammatory properties (Chi et al., 2023; Galbraith et al., 2023; Rachubinski et al., 2024; Sullivan et al., 2017; Sullivan et al., 2016; Tuttle et al., 2020; Waugh et al., 2023). Furthermore, fasudil treatment upregulated genes involved in SHH signaling, which is essential for proper neurogenesis throughout the developing brain (Belgacem et al., 2016). Previous studies in the Ts65Dn mouse model of DS revealed deficits in SHH signaling (Gao et al., 2021).

Furthermore, restoring the SHH pathway in the early prenatal period improved cellular proliferation in the cerebellum and corrected behavioral deficits in learning / memory and hyperactivity (Das et al., 2013; Gao et al., 2021; Roper et al., 2006). Fasudil’s amelioration of SHH pathway gene dysregulation suggests that fasudil administration during early neuronal development might also improve these cell growth and behavioral phenotypes. Overall, the results of this study demonstrate fasudil’s potential to simultaneously target multiple T21-associated abnormalities in a beneficial manner.

Our *in vitro* results using human T21 NPCs complement those of an *in vivo* fasudil study using the Ts65Dn mouse model of DS (Lopez-Hidalgo et al., 2024). Chronic treatment of Ts65Dn mice with fasudil from postnatal day (P) 0 – P28 improved cell proliferation and neurogenesis of the subgranular zone of the hippocampus, corrected abnormal dendritic morphology of trisomic hippocampal neurons, and reduced inhibitory synapses, which are more numerous in both Ts65Dn and T21 (Lopez-Hidalgo et al., 2024). Thus, different preclinical model systems and outcome measures both point to the potential therapeutic effects of fasudil on neurological development.

Fasudil inhibits both ROCK1 and ROCK2, affecting multiple ROCK downstream targets. Therefore, fasudil and other ROCK inhibitors have the potential to alleviate numerous pathologies, including neurodegeneration, cancer, glaucoma, and kidney failure (Cheng et al., 2025; Feng et al., 2016). Fasudil has been used clinically in Japan since 1995 to prevent cerebral vasospasm after subarachnoid hemorrhage (Roskoski, 2023; Shibuya et al., 1992; Zhao et al., 2006). The beneficial effects of fasudil in preclinical models of neurodegenerative disorders (Koch et al., 2018; Song et al., 2013; Takata et al., 2013; Yang et al., 2020), which may relate to its ability to block immune system / microglial activation (Roser et al., 2017; Wang et al., 2022), have led to fasudil’s employment in multiple clinical trials related to neurodegeneration (clinicaltrials.gov) (Koch et al., 2024; Wolff, Bidner, et al., 2024; Wolff, Peine, et al., 2024). Of note, fasudil shows inhibitory effects on multiple protein kinase pathways in addition to ROCK1 and ROCK2 (https://www.guidetopharmacology.org/GRAC/LigandScreenDisplayForward?ligandId=51 <u>81CscreenId=3</u>). This concern is being addressed by the development of more selective analogs such as ripasudil (Cheng et al., 2025; Feng et al., 2016; Koch et al., 2018; Ye et al., 2024).

ROCKs perform essential functions during earlier embryonic developmental stages and organogenesis (Shi & Wei, 2022), and thus application of fasudil at these early time points could be detrimental. However, animal studies suggest the feasibility of prenatal administration of fasudil during later gestational stages, which would align with human fetal stages that occur after diagnosis of T21. In rats, fetal exposure to fasudil reduced pre- eclampsia (Gu et al., 2017) and reduced intrauterine growth restriction, although this was also associated with greater postnatal body weight, increased food intake, and metabolic changes (Butruille et al., 2012). Fasudil also reduced the myogenic (vasoconstriction) response in acute ductus arteriosus compression in late gestation fetal sheep (Tourneux et al., 2008).

Anti-inflammatory properties have been previously reported for fasudil (Guo et al., 2020; Hou et al., 2012; Liu et al., 2022; Okamoto et al., 2010; Song et al., 2013), EGCG (Capasso et al., 2025), apigenin (Guedj et al., 2020), and downstream metabolic products of sphingosine (Gomez-Larrauri et al., 2025). To the best of our knowledge, the anti-inflammatory effects of isoxsuprine described in this study are novel. While anti-inflammatory effects of fasudil were identified in T21 NPCs as well as LPS-challenged microglia systems, isoxsuprine only showed inflammatory effects in the microglia studies. The different anti-inflammatory properties of candidate pharmacotherapies may reflect cell-specific or developmental stage-specific effects of each molecule that will need to be elucidated further. Microglia may play important roles in T21 CNS phenotypes (Flores-Aguilar et al., 2020; Jin et al., 2022; Martini et al., 2020; Pinto et al., 2020; Tiberi et al., 2025), so additional studies on trisomic microglia would be warranted.

Although EGCG and sphingosine showed fewer T21 DEGs in comparison to fasudil, most gene changes for EGCG and sphingosine were predicted to worsen T21-associated gene dysregulation, suggesting these molecules would not benefit neurodevelopment and could potentially cause harm. This is of interest because some parents administer dietary supplements such as EGCG/green tea extract to their children with DS, often without informing medical caregivers (Lewanda et al., 2018). EGCG studies have shown mixed results in individuals with T21 and mouse models of DS. While several studies report beneficial effects on skeletal/craniofacial, cognitive, behavioral, and cardiac phenotypes (Blazek et al., 2015; Catuara-Solarz et al., 2015; de la Torre et al., 2016; De la Torre et al., 2014; McElyea et al., 2016; Souchet et al., 2019; Stagni et al., 2016; Starbuck et al., 2021; Tielemans et al., 2025), other studies showed no effects (Cieuta-Walti et al., 2022), deleterious effects (Goodlett et al., 2020; Tielemans et al., 2025), or variable effects related to dosage, timing of administration, or type of commercially available supplement (Abeysekera et al., 2016; Jamal et al., 2022; Stringer et al., 2017). Controlled preclinical and clinical studies are needed to understand the complex effects of these molecules and thus allow clinicians to advise parents in an informed manner.

The transcriptomic-based approach used in this study did not identify new molecules with beneficial effects on cell growth or inflammation in NPCs. While this may reflect limitations of the testing used in our NPC model system, it may also relate to distinct transcriptional responses of Eup and T21 cells to the same molecule. Eup and T21 NPCs showed strikingly different gene expression responses for all five pharmacotherapies examined by RNA-Seq in this study. This suggests that T21 cells may respond to candidate therapeutics in ways that cannot be predicted from Eup cells, and thus Eup cell-derived gene signatures may not accurately predict the pharmacological effects on transcription in T21 cells. Future queries of LINCS with targeted subsets of T21 DEGs – for example, gene signatures that only include inflammatory or oxidative stress pathways – may be more informative than queries using the full gene expression signature. Newer deep learning methods for predicting treatment responses that could be trained on these new data from T21 NPCs might also improve cell-specific connectivity results. In addition, the negative effects on cell growth for eight candidate molecules (Supp. Tab. S1) could have been caused by targeting T21 gene dysregulation that promotes survival in the context of aneuploidy. If true, these essential T21 pathways will need to be defined in future studies so that they can remain unchanged during application of pharmacological treatments.

In summary, we demonstrated that euploid and T21 NPCs showed distinct transcriptomic responses to five candidate pharmacotherapies. We also determined that the ROCK inhibitor fasudil improved growth in T21 NPCs, normalized expression of inflammatory genes in T21 NPCs, and reduced inflammatory markers in LPS-stimulated microglia model systems. Our results, combined with those from a prior *in vivo* study (Lopez-Hidalgo et al., 2024), suggest fasudil merits serious consideration for further study as a potential prenatal pharmacotherapy.

## Materials and methods

### Selection of candidate pharmacotherapies using LINCS

To identify novel therapeutic candidate molecules via a transcriptomics approach, connectivity mapping was performed on a connectivity database created from a subset of the 2017 LINCS GEO releases, reflecting 80 cell types and 1330 named molecules, previously described as the larger "sparse" matrix (Sapashnik, et al., 2023). This data set was then completed by imputing missing data with the neighborhood collaborative filtering method, as described in Section 3.5 of Sapashnik et al., 2023, to provide data for any cell/molecule combinations not experimentally evaluated in the LINCS data. The query signature dataset contained 76 upregulated and 58 downregulated genes (Supp. Tab. S1e), which corresponded to all genes within the T21 NPC DEG list (FDR-adjusted P-value 0.1, FC ≥|1.2|) previously published by our lab (Lee et al., 2025) that are also represented on the L1000 platform.

The 165 molecules with LINCS scores <-0.2 in NPCs were screened for toxicity and teratogenicity by searching PubMed for published articles on each molecule using the molecule name and toxic or teratogenic as search terms. Molecules with reported teratogenicity or contraindications in pregnancy were eliminated from consideration for further testing, and only molecules with no or limited toxic effects and limited or mild adverse effects were used for testing. A subset of 17 molecules from this list was chosen for further examination using NPC cultures.

### Cell culture

Nine Eup and eight T21 iPSC lines previously generated in our laboratory (Lee et al., 2025) were used for these experiments (Supp. Tab. S2). iPSC lines were stably transduced with Nuclight Red Lenti, EF1a, puro (4625, Sartorius, Gottingen, Germany) per manufacturer’s instructions, which generated a nuclear-restricted fluorescent signal that enabled live cell imaging. Stable selection for the Nuclight Red vector was performed using 1 µg/ml puromycin and subsequent maintenance was performed using 0.5 µg/ml puromycin. iPSCs were cultured on Corning Matrigel-coated plates (354277, Fisher Scientific, Waltham, MA) in Gibco StemFlex media (A3349401, ThermoFisher Scientific, Waltham, MA), and colonies were passaged as aggregates at 60-70% confluency using ReLeSR (100-0483, StemCell Technologies, Vancouver, BC).

Neural induction of stably transduced iPSCs to generate NPCs was performed as previously described (Lee et al., 2025). Briefly, Nuclight Red-transduced Eup and T21 iPSC lines were dissociated to single cells at passage ten or greater following transduction using Accutase (07920, Stem Cell Technologies) and plated at 1 X 10^5^ cells/cm^2^ on Matrigel-coated plates using the STEMdiff SMADi Neural Induction Kit (08581, StemCell Technologies). All subsequent passages used Accutase. After culturing for three passages in StemDiff Neural Induction Media, cells were switched to STEMdiff Neural Progenitor Media (05833, StemCell Technologies), and lines were subsequently maintained on Matrigel in this media. After three passages, NPCs were cryopreserved in Cryostor CS10 (100-1061, StemCell Technologies) at each passage up to passage 10. All NPC assays were performed between passages three and ten, with most assays performed between passages five and seven.

At each passage during neural induction and NPC maintenance, media was supplemented with CEPT cocktail (Chen et al., 2021) at the following final concentrations: 50 nM chroman 1 (7163, Bio-Techne, Minneapolis, MN), 5 µM emricasan (S7775, Selleck Chemicals), 1:1000 dilution of polyamines (P8483, MilliPoreSigma), and 0.7 µM trans-ISRIB (5284, Bio-Techne). Chroman 1 is a ROCK inhibitor, emricisan is a pan-caspase inhibitor, polyamines support cell growth and cell attachment, and trans-ISRIB selectively inhibits the integrated stress response. Media containing CEPT was removed within 24 hrs of passaging and replaced with unsupplemented media.

Maintenance of the expected T21 karyotype in trisomic lines along with the absence of additional chromosomal abnormalities along was confirmed in NPCs by Karyostat analyses (ThermoFisher Scientific). NPC cultures were confirmed to be free of mycoplasma contamination by PCR testing of media using a mycoplasma PCR detection kit (G238, Applied Biological Materials, Richmond, BC, Canada).

### Analysis of the effects of CEPT and Y-27C32 on growth

Three Eup lines (EU6, EU7, EU12) and 3 T21 iPSC lines (TS3, TS4, TS8) were cultured in 6-well plates and subjected to neural induction, as described above. One cell line was plated per plate at 1 X 10^5^ cells/cm^2^ at each passage. Three wells were passaged with media containing CEPT and the other three with media containing 10 µM Y-27632. Cells were cultured for three passages, and growth from passage zero through passage three was monitored by live-cell imaging of fluorescent nuclei at 594 nM every 24 hrs. Imaging was performed using a Cytation 5 Cell Imaging Multimode Reader (Agilent Technologies, Santa Clara, CA), and cell counts were measured using Gen5 version 3.12 imaging software (Agilent Technologies), with object sum area of fluorescent nuclei values used to measure cell number. Ratios of CEPT-treated cell counts to Y-27632 cell counts were calculated for each cell line at 1, 2, and 3 days after each passage.

### Quantitation of NPC growth in the presence of candidate pharmacotherapies

Eup and T21 NPCs were plated with NPC media containing CEPT at an initial density of 5000 cells/well (1.0E+05 cells/ml) in 384-well black optical bottom plates (164586, ThermoFisher Scientific) or 75,000 cells/well (3.75E+04 cells/ml) in standard 6-well tissue culture plates. The 6-well plate studies were performed after the 384-well plate studies, to examine molecule effects in a larger well format in which more consistent cell growth could be achieved over longer time periods and sufficient RNA could be collected for RNA-Seq.

NPCs were cultured for 120 hrs in the presence or absence of candidate pharmacotherapeutic molecules. At 12 hrs, media was changed to remove CEPT and was replaced with either NPC media containing candidate molecules (treated wells) or unsupplemented NPC media (untreated wells). This media change was repeated 48 hrs later (at 60 hrs). Two or three concentrations of each molecule were used, based on previous *in vitro* dose range-finding studies of each molecule (Supp. Tab. 1). Each molecule was tested in two independent plates/experiments in 4 replicate wells per line/condition for the 384-well plate format, and in 6 replicate wells per line/condition for the 6-well plate format. Quantitative imaging was performed using a Cytation 5 Cell Imaging Multimode Reader (Agilent Technologies) and Biospa 8 Automated incubator (Agilent Technologies) system along with Gen5 version 3.12 software (Agilent Technologies) to calculate the object sum area of fluorescent nuclei every 12 hrs for 384-well plate cultures and every 24 hrs for 6-well plate cultures. Subtractive normalization was performed on treated wells using the untreated well object sum area values in the same experiment/plate for each NPC line.

Doubling time (DT) was calculated by the equation DT = ln(2) / k, in which the growth rate (k) was calculated by the polyfit function in Python using the least square method and the general equation for cell growth (N(t) = (N0) * kt, where N = cell number/object sum area at specified time point, N0 = initial cell number/object sum area, t = time). Outliers were removed by: 1) not meeting a minimum threshold of continuous growth for ≥ 48 hrs, 2) by using the interquartile range (IQR) outlier formula for DT for each set of 4 or 6 replicates (IQR = q3 – q1; lower limit = q1 – 1.5 * IQR, upper limit = q3 +1.5 * IQR), or 3) not meeting a minimum data point count of 3 for each set of 4 or 6 replicates (for 384-well or 6 well experiments, respectively) following the previous two outlier removal steps. DT calculations and outlier analyses were done using a Jupyter notebook written in Python. Within-plate comparisons of DTs for untreated and treated wells of the same cell line were tested by Mann-Whitney U test in Python and in GraphPad Prism, with a significance threshold of p < 0.05. Two-way ANOVA analyses of growth curves were performed using GraphPad Prism.

### RAW 2C4.7 cell culture and inflammation assays

The mouse monocyte/macrophage-like cell line RAW 264.7 was purchased from ATCC (TIB-71), and cultured following ATCC recommendations in DMEM + 10% FBS (10569-010 and 16140-071, Thermo Fisher Scientific). Compounds were assessed in two studies: (1) treatment with no LPS to evaluate compound tolerability / compound-induced cytotoxicity (1, 10 and 100 µM); (2) treatment at 1, 10 and 30 µM followed by exposure to LPS (10 ng/ml) to mitigate LPS-induced inflammation. Cells were plated in 24-well plates at a density of either 187 K (study 1) or 240 K cells per well (study 2). The next day, cells were pre-treated with fasudil, apigenin, EGCG, isoxsuprine HCl, and sphingosine using the vehicle dimethyl sulfoxide (DMSO, Sigma, Cat # D2650), and control cells were pre-treated with the same dilution of DMSO. One hr following the addition of compound/vehicle, the cells were challenged with 10 ng/ml LPS (L4005, Sigma-Aldrich, St. Louis, MO, USA; *Escherichia coli* O55:B5), a dose that induced a substantial and sub-maximal inflammatory response in prior pilot studies (Tweedie et al., 2011; Tweedie et al., 2009).

After 20-24 hrs, media was collected and used for downstream assays. Cell viability was determined using the CytoTox-ONE Homogeneous Integrity Assay (G7890, Promega), culture media levels of nitrite were assessed using the Nitrate/Nitrite Fluorometric Assay Kit (KA1344, ABNOVA), and culture media levels of cytokines TNF-α and IL-6 were determined by Enzyme-linked ImmunoSorbent Assays (ELISAs): ELISA MAX™ Deluxe Set Mouse TNF-α (430915, BioLegend Inc) and ELISA MAX™ Deluxe Set Mouse IL-6, (431315, BioLegend Inc). All assays were performed as directed by the manufacturers. Data were normalized to the control LPS+DMSO group, i.e., inflammation without drug treatment, and then expressed as percent change from the control cells, with control data defined as 100%. Data are presented as mean± S.E.M. of N observations (N = 3, except for apigenin IL-6, 30 µM, where N = 1). Data were processed by GraphPad Prism, version 10.0.3. Values were assessed for outliers (ROUT) and normal distributions (Shapiro-Wilk test). Treatment and control values were compared using ordinary one-way ANOVA followed by Dunnett’s multiple comparisons test, or Kruskal-Wallis test ANOVA followed by Dunn’s multiple comparisons test.

### IMG cell culture and inflammation assays, and immunocytochemistry (ICC)

The immortalized adherent mouse microglial “IMG” cell line (SCC-134, Sigma-Aldrich; RRID:CVCL_HC49) was cultured in High Glucose DMEM (D6546, Sigma) with 10% heat-inactivated fetal bovine serum (10082147, Gibco), 1× L-Glutamine (TMS-002-C, Sigma) and 100 U/mL penicillin/streptomycin (15140148, Gibco). IMG cell culturing was performed according to the product datasheet, with cell passaging performed with Accutase (A6964, Sigma). Cells were used for a maximum of 10 passages.

IMG cells for cytokine assays were grown in 24-well (40 K cells/well) tissue culture-treated plates in 500 μl of media. Cells used for immunocytochemistry were plated in 96-well sterile glass bottom plates (P69-1.5H-N, Cellvis) at a density of 30,000 cells/well and were grown to ∼80% confluency (1–2 days). Fresh media was added on day 1 after plating and changed every two days, as needed. To assess for possible toxicity induced by fasudil HCl (S1573, Selleck Chemicals), initially IMG cells were cultured in the presence of fasudil alone (no LPS) in 24 well plates in 500 µl of basic growth media. The following day, cells were treated with fasudil prepared in the same cell culture media (vehicle), at 1, 10, 30, 50 or 100 μM. After 24 hrs, the media was collected and replaced with 500 μl fresh media to allow assessment of cell viability. Fasudil was toxic to IMG cells at 100 μM (compared to vehicle control), thus 50 µM was the highest concentration used in LPS-induced inflammation related studies. IMG cell viability was quantified using the CellTiter 96® Aqueous One Solution Cell Proliferation Assay kit (MTS) (G3580, Promega, Madison, WI, USA) according to the manufacturer’s protocol. IMG cells were pretreated with fasudil at 1, 10, 30, or 50 μM and then, one hr later, challenged with LPS at 10 ng/mL in the continued presence of fasudil. Following a 24 hr incubation, conditioned media was collected and replaced with 500 μl of fresh, basic growth media to allow the assessment of cell viability. Conditioned media was later analyzed for TNF-α levels by ELISA using the Mouse TNF-α ELISA MAX™ Deluxe Set (430904, BioLegend), according to manufacturer recommendations. Data are presented as mean± S.E.M. of n observations (N = 3-4). Data were processed by GraphPad Prism and statistical analysis involved ordinary one-way analysis of variance (ANOVA) with Dunnett’s comparison (vs. vehicle or 0 μM fasudil).

For immunocytochemistry (ICC), experimental conditions were replicated in triplicate. Following 1 hr pretreatment with fasudil (50 µM), isoxsuprine HCl (50 µM) (S5669, Selleck Chemicals), or vehicle (media), 10 x LPS (final concentration 10 ng/mL) was added for 15 minutes. Cells were then washed twice with ice-cold PBS and fixed with cold 4% PFA (AAJ19943K2, Fisher Scientific) for 10 min. PFA was removed and cells were washed three times with cold PBS and stored at 4°C in PBS prior to staining. Cells were permeabilized and blocked in microglia staining buffer (MSB) (3% BSA and 0.1% saponin (47036, Sigma) in PBS) for 1 hour at room temperature (RT), followed by incubation with the primary antibody anti-NF-κB (6956S, Cell Signaling, dilution 1:500, raised in mouse) in MSB for 1 hour at RT. After primary antibody incubation, cells were washed three times with 0.1% saponin in PBS (wash buffer) for 5 min/wash. The corresponding highly cross-adsorbed Alexa Fluor™ secondary antibody goat anti-mouse 555 IgG (A-21422, Invitrogen, RRID:AB_2535844) was diluted (1:500) in MSB and applied to cells for 1 hour at RT. Primary Phalloidin staining (A12379, Invitrogen, 488 fluorophore, dilution 1:500) was performed with secondary antibody incubation. Following secondary antibody staining, cells were stained with 4’,6-Diamidino-2-Phenylindole, Dilactate (DAPI) (D3571, Invitrogen, dilution 1:1000) for 10 min to visualize cell nuclei (at 405nm wavelength of excitation). Cells were washed 3 X 5 mins in wash buffer with a fourth wash in PBS alone. Cells were stored in 100 µl of PBS for imaging. Cells were imaged at 20 X using a Nikon Eclipse Ti2 confocal microscope. A Z-stack was collected through each cell culture and collapsed into a maximum intensity projection for the field of view (FOV) (1024 µm x 1024 µm). A representative single FOV is shown for each experimental group. Imaging parameters were calibrated using secondary-only controls to set laser power and gain.

### RNA isolation and RNA-Seq

Eup and T21 NPCs that had been cultured in the 6-well plate format were used for RNA collection. On the final day of culture, media was removed and 450 µl TriReagent from Direct-zol MiniPrep kits (R2053-A, Zymo Research, Irvine, CA) was added to each well. A cell scraper was used to make a homogeneous lysate, then samples were pipetted into Eppendorf tubes and stored at -80°C in Tri reagent until RNA collection. Total RNA was subsequently collected from 3 untreated and 3 treated wells for each NPC line using Direct-zol MiniPrep kits, following the manufacturer’s instructions. RNA concentration and purity was determined using a Nanodrop 8000 (Thermofisher), and RNA integrity (RIN) scores were generated using a Bioanalyzer (Agilent). Most RNA samples had RIN scores ≥9, and only samples with RIN scores ≥8.4 were used for sequencing.

1000 ng of total RNA for each sample were used for RNA-Seq. Strand-specific, poly-A selected RNA-Seq libraries were generated using NEBNext Ultra™ II Directional RNA Library Prep Kits for Illumina and NEBNext Poly(A) mRNA Magnetic Isolation Modules (New England Biolabs). Each library was tagged with unique dual indexes. Sequencing of the pooled libraries was performed at the NIH Intramural Sequencing Center (NISC) on a NovaSeqX Plus Sequencing System (Illumina) on 25B flow cells. At least 40 million 150-base read pairs were generated for each individual library. RNA-Seq data files and associated metadata have been submitted to dbGaP, study phs004455.

### RNA-Seq analyses

Data were analyzed by ROSALIND (San Diego, CA, https://rosalind.bio/). Reads were trimmed using cutadapt1. Quality scores were assessed using FastQC2. Reads were aligned to GRCh38 using STAR3. Individual sample reads were quantified with HTseq4 and normalized via Relative Log Expression (RLE) using DESeq2 R library5. Sample MDS plots were generated as part of the QC step using RSeQC6. DESeq2 was used to calculate fold changes and p-values and perform optional covariate correction. Clustering of genes for the final heatmap of differentially expressed genes was done using the Partitioning Around Medoids (PAM) method using the fpc R library7. Samples that did not pass quality control were removed prior to differential gene expression analysis. Four Eup and 3-4 T21 NPCs lines were used for each RNA-Seq analysis.

Cell line was used for covariate correction for T21 only and Eup only analyses, to remove confounding differences among cell lines within each genotype. Sex was used for covariate correction for T21Unt and EupUnt comparisons, since male and female lines showed separation by MDS. Each candidate molecule was tested independently, thus T21Unt vs EupUnt DEG datasets were generated for each candidate molecule analysis. To determine if candidate molecules corrected or worsened T21-associated gene dysregulation, the DEG list generated from the same experiment was compared with candidate molecule-induced DEGs.

Pathway analyses were performed using the ROSALIND knowledgebase, Metascape 3.5 (Zhou et al., 2019), or Ingenuity Pathway Analysis (IPA, Winter 2025 release, Qiagen).

## Acknowledgments

This research was supported in part by the Intramural Research Program of the National Institutes of Health (NIH) (1ZIA HG200399 and AG000311) and by NSF award CCF-1934553 (D.Z and D.K.S.). We thank the NIH Intramural Sequencing Center for library construction and sequencing for bulk RNA-Seq analyses and Nicole Briceno for critical reading of the manuscript. The contributions of the NIH authors are considered Works of the United States Government. The findings and conclusions presented in this paper are those of the authors and do not necessarily reflect the views of the NIH or the U.S. Department of Health and Human Services.

---

**Supplemental figure S1.**
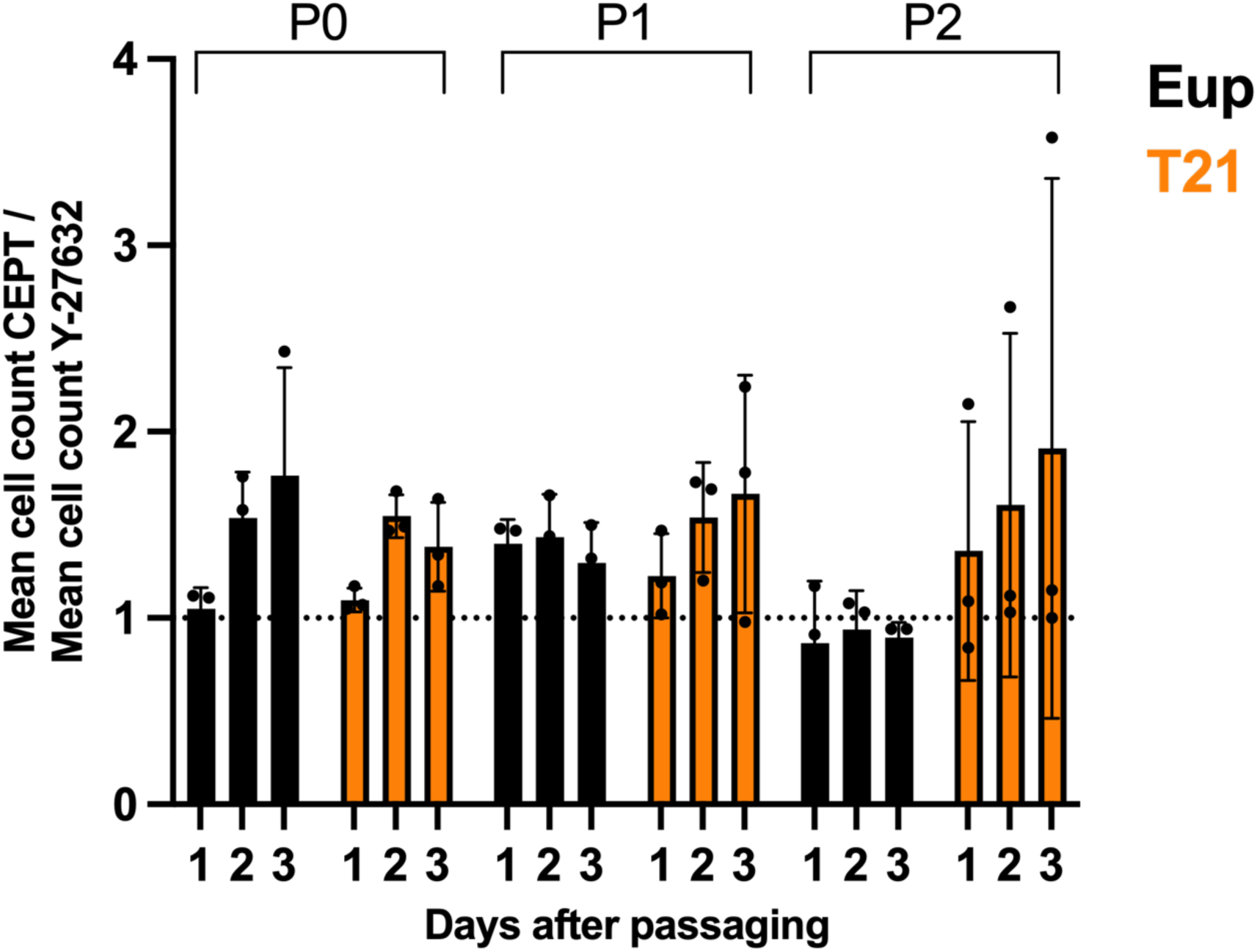
Passaging with CEPT improved growth during neural induction of NPCs from iPSCs. More cells survived passaging with CEPT supplementation in comparison to Y-27632 supplementation in Eup and T21 lines. Ratios of CEPT-treated cell count to Y-27632 cell count in the same line are shown at one, two, and three days after passages 0, 1, and 2 (P0, P1, and P2, respectively). Ratio values >1 indicate greater cell numbers in CEPT-treated relative to Y-27632-treated lines. N = 3 Eup lines (EU6, EU7, EU12) and 3 T21 lines (TS3, TS4, TS8). Each data point = the mean of 3 replicates. Black = Eup lines, orange = T21 lines.

**Supplemental figure S2.**
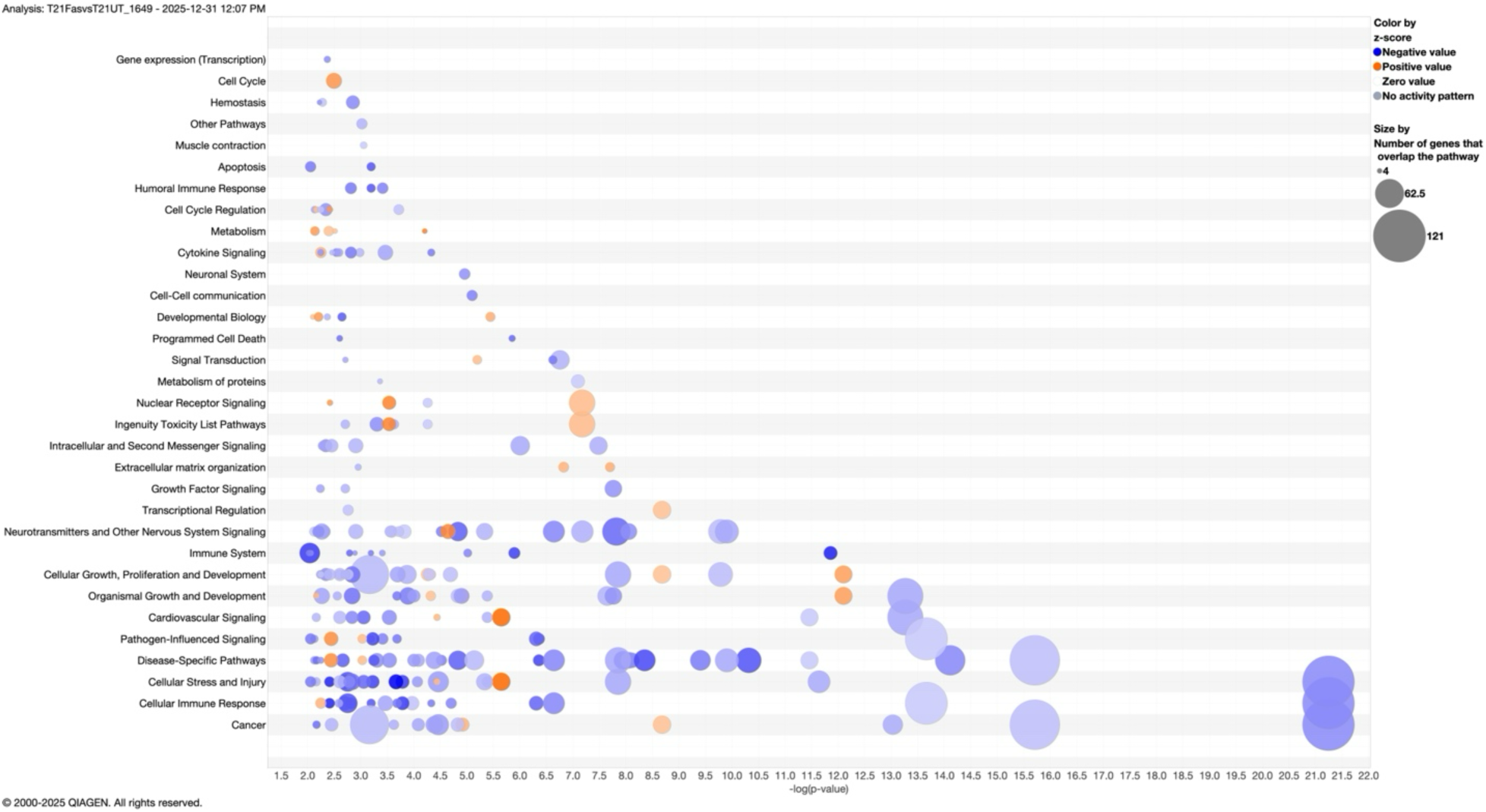
Significantly enriched pathways in fasudil-treated T21 NPCs. Ingenuity Pathway Analysis for the 1649 T21 DEGs showed categories of significantly enriched pathways, including downregulation of numerous cellular immune response / immune system and cellular stress and injury pathways. Blue = negative Z score / pathway downregulation, and orange = positive Z score / pathway upregulation. Circle size represents the number of DEGs in the pathway.

## Supplemental Tables

**Supplemental Table S1.** Candidate pharmacotherapies tested for effects on growth of euploid and T21 NPCs.

**Supplemental Table S2.** Euploid and T21 iPSCs and NPCs used in this study.

**Supplemental Table S3**. Viability and inflammatory marker results in LPS-challenged RAW 264.7 cells and IMG cells.

**Supplemental Table S4**. Comparison of gene expression in untreated T21 vs. untreated euploid NPCs.

**Supplemental Table S5**. Comparison of gene expression in fasudil-treated T21 vs. untreated T21 NPCs.

**Supplemental Table S6**. Identification of 1356 overlapping DEGs between T21Fas DEGs and T21 DEGs.

**Supplemental Table S7**. Comparison of gene expression in fasudil-treated euploid vs. untreated euploid NPCs.

**Supplemental Table S8.** Comparison of gene expression in EGCG-treated T21 vs. untreated T21 NPCs.

**Supplemental Table S9.** Comparison of gene expression in sphingosine-treated T21 vs. untreated T21 NPCs.

**Supplemental Table S10.** Comparison of gene expression in apigenin-treated T21 vs. untreated T21 NPCs and isoxsuprine-treated T21 vs. untreated T21 NPCs.

**Supplemental Table S11.** Comparison of gene expression in EGCG-treated, apigenin-treated, sphingosine-treated, and isoxuprine-treated euploid NPCs vs. untreated euploid NPCs.

